# Optimized hip-knee-ankle exoskeleton assistance at a range of walking speeds

**DOI:** 10.1101/2021.03.26.437212

**Authors:** Gwendolyn M. Bryan, Patrick W. Franks, Seungmoon Song, Alexandra S. Voloshina, Ricardo Reyes, Meghan P. O’Donovan, Karen N. Gregorczyk, Steven H. Collins

**Affiliations:** Department of Mechanical Engineering, Stanford University, Stanford, USA; Mechanical and Aerospace Engineering, University of California, Irvine, Irvine, USA; U.S. Army Natick Soldier Research, Development and Engineering Center, USA

**Keywords:** Exoskeleton, Augmentation, Walking speed, Human-in-the-Loop Optimization

## Abstract

**Background:** Effective autonomous exoskeletons will need to be useful at a variety of walking speeds, but we do not know how optimal exoskeleton assistance should change with speed. Optimal exoskeleton assistance may increase with speed similar to biological torque changes or a well-tuned assistance profile may be effective at a variety of speeds.

**Methods:** We optimized hip-knee-ankle exoskeleton assistance to reduce metabolic cost for three participants walking at 1.0 m/s, 1.25 m/s and 1.5 m/s. We measured metabolic cost, muscle activity, exoskeleton assistance and kinematics. We performed two tailed paired t-tests to determine significance.

**Results:** Exoskeleton assistance reduced the metabolic cost of walking compared to wearing the exoskeleton with no torque applied by 26%, 47% and 50% at 1.0, 1.25 and 1.5 m/s, respectively. For all three participants, optimized exoskeleton ankle torque was the smallest for slow walking, while hip and knee torque changed slightly with speed in ways that varied across participants. Total applied positive power increased with speed for all three participants, largely due to increased joint velocities, which consistently increased with speed.

**Conclusions:** Exoskeleton assistance is effective at a range of speeds and is most effective at medium and fast walking speeds. Exoskeleton assistance was less effective for slow walking, which may explain the limited success in reducing metabolic cost for patient populations through exoskeleton assistance. Exoskeleton designers may have more success when targeting activities and groups with faster walking speeds. Speed-related changes in optimized exoskeleton assistance varied by participant, indicating either the benefit of participant-specific tuning or that a wide variety of torque profiles are similarly effective.

## 1 Introduction

We expect that autonomous exoskeletons will need to be effective at a variety of walking speeds. These devices have the potential to assist a wide range of people, from those with walking impairments to military personnel. These populations walk at different self-selected speeds; for example, able-bodied adults walk between 0.75 and 1.75 m/s [1, 2], while adults who have had a stroke walk between 0.08 and 1.05 m/s [3]. To effectively assist people during natural walking conditions, exoskeletons will need to be useful at a variety of walking speeds.

Exoskeletons have successfully reduced the metabolic cost of walking when optimized at a single speed. Typically, they have assisted able-bodied participants walking between1.25 and 1.4 m/s [4]. Some studies have assisted able-bodied participants walking as slowly as 1.1 m/s [5] and others have assisted able-bodied participants walking as quickly as 1.5 m/s [6–8]. Many successful exoskeletons apply torque at the hip, ankle or both [5–15] and have reduced the metabolic cost of walking by up to 24% [12]. Whole-leg assistance can reduce metabolic cost even further. Hip-knee-ankle exoskeleton assistance has reduced the metabolic cost of walking at 1.25 m/s by 50% [16].These effective assistance strategies are well-tuned at one speed, but future exoskeleton products will need to be effective at a variety of speeds

Optimized assistance at a range of speeds could inform hardware requirements and assistance changes in future devices. One study determined the optimal spring stiffness for spring-like ankle exoskeleton assistance while participants walked at 1.25, 1.5 and 1.75 m/s [17]. The same spring stiffness produced the largest metabolic reductions for both 1.25 and 1.75 m/s, suggesting that consistent assistance may be effective at a range of speeds. A pilot study with bilateral ankle exoskeletons optimized exoskeleton assistance at 0.75, 1.25 and 1.75 m/s [12] which resulted in reductions to metabolic cost of 3%, 33% and 39%, respectively. In that study, ankle torque optimized to zero for the slow walking speed, suggesting no benefit was possible through exoskeleton assistance, and increased from the medium to fast walking speeds, suggesting higher torques may be optimal at higher speeds. Further investigation of optimized torque changes with speed could determine whether assistance should change with speed and, if so, how it should change, informing the design of future devices.

Here, we optimized whole-leg exoskeleton assistance for different walking speeds to minimize the metabolic cost of walking. We hypothesized that exoskeleton assistance would result in lower metabolic rate at all speeds, but with less benefit at slower speeds, and that optimized exoskeleton assistance would increase with speed. We used human-in-the-loop optimization of assistance applied by a hip-knee-ankle exoskeleton emulator [18] while participants walked on a treadmill at 1.0, 1.25 and 1.5 m/s, representative of typical self-selected speeds [1]. We evaluated changes in metabolic cost, muscle activity, exoskeleton assistance and kinematics across speeds and across assistance conditions within each speed. We used biomechanics measurements to gain insights into the mechanisms underlying any trends in metabolic rate. We expected results from this study to be used to prescribe effective speed-related assistance changes for future exoskeleton products.

## 2 Methods

We optimized hip-knee-ankle exoskeleton assistance at slow, medium, and fast walking speeds (1.0 m/s, 1.25 m/s, and 1.5 m/s). We used human-in-the-loop optimization to minimize the metabolic cost of walking at each speed. Human-in-the-loop optimization is a strategy that optimizes exoskeleton assistance in response to real-time participant performance metrics [12].

Three able-bodied participants (1F 2M, age 26-36 years, 60-90 kg, 170-188 cm, expert users) wore a hip-knee-ankle exoskeleton emulator [18] while walking on a split-belt treadmill. The exoskeleton can apply bilateral assistance in hip flexion and extension, knee flexion and extension, and ankle plantarflexion. All participants were expert users; two had participated in previous experiments and the third underwent a lengthy training protocol prior to beginning the optimization (Supplementary Section 1). All experiments were approved by the Stanford University Institutional Review Board and the US Army Medical Research and Materiel Command (USAMRMC) Office of Research Protections. Participants provided written and informed consent.

### 2.1 Exoskeleton Hardware

The hip-knee-ankle exoskeleton emulator can apply torque about the joints using powerful, offboard motors and a Bowden cable transmission ([18]; Figure 1). The end-effector has a worn mass of 13.5 kg, and was fit to participants through length and width adjustability and interchangeable boots.

**Figure 1.**
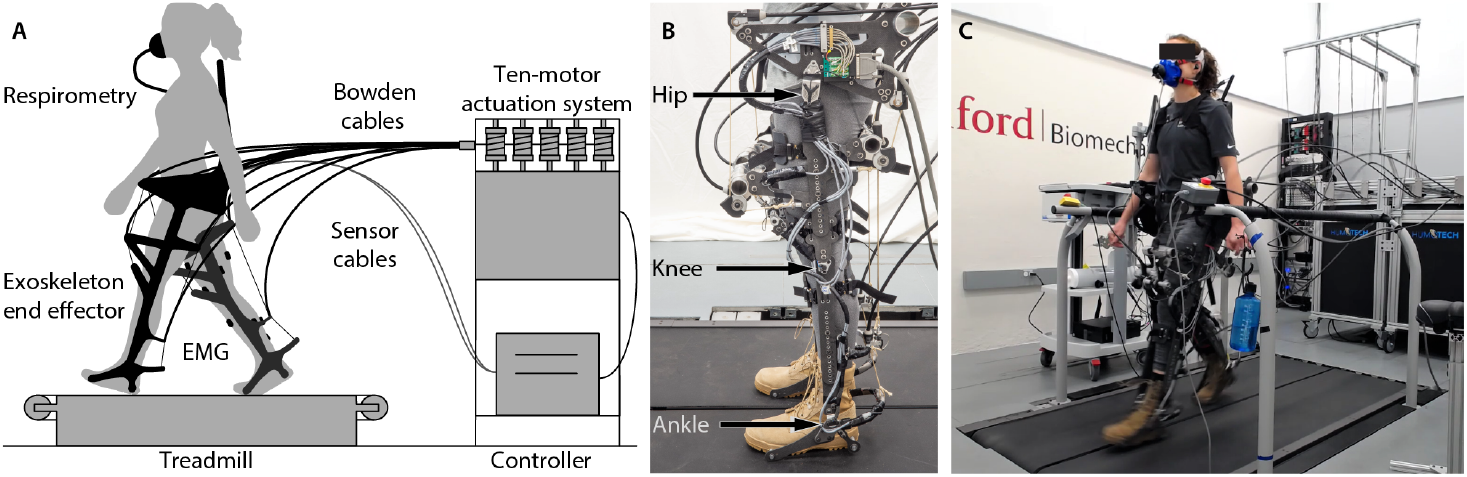
Experimental setup. (A) exoskeleton emulator system. (A) exoskeleton emulator system. A participant wears the hip-knee-ankle exoskeleton emulator and walks on a split-belt treadmill. Powerful, offboard motors apply joint torques through a Bowden cable transmission. (B) exoskeleton end effector. The device can assist the hips, knees and ankles. (C) experimental setup. Metabolic rate and muscle activity are measured while a participant wears the exoskeleton end effector and walks on a split-belt treadmill.

### 2.2 Exoskeleton Control

We applied hip, knee, and ankle exoskeleton assistance profiles as a function of stride time [18]. These profiles are defined by 22 parameters that can be adjusted by the optimizer (Figure 2). The torque parameterization was successful in a previous optimization study with this device [16], and was inspired by biological torque data, effective single joint assistance and pilot experiments with different parameterization strategies. The profiles were generated with splines, and parameter ranges were limited for participant comfort (Supplementary Section 8).

The hip, knee and ankle profiles were defined by a total of 22 parameters. The hip profile, defined by eight parameters, applied hip extension torque during early stance and hip flexion torque during swing (Figure 2 Hips). Knee assistance applied a virtual spring during early stance, knee flexion torque near toe-off and a virtual damper during swing for a total of ten tunable parameters (Figure 2 Knees). Ankle assistance applied plantarflexion torque near toe-off and was defined by four parameters (Figure 2 Ankles). A more detailed description can be found in the supplementary materials (Supplementary Section 2).

**Figure 2.**
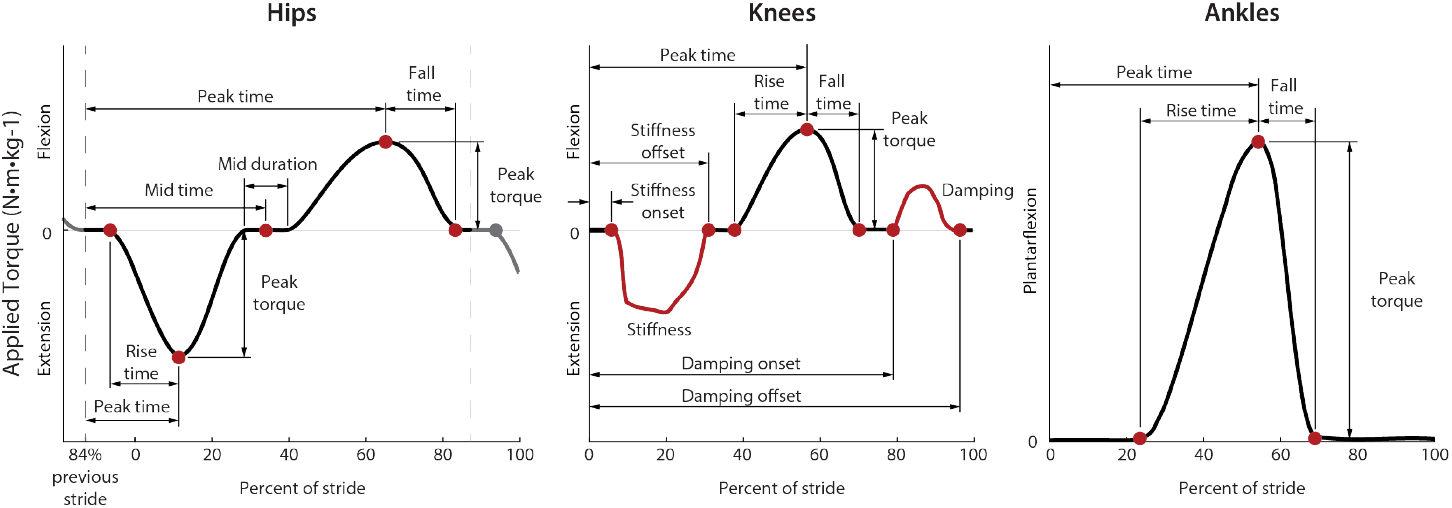
Parameterization of the hip, knee and ankle exoskeleton profiles. Assistance profiles defined torque as a function of stride time with periods of state-based torque at the knees. Hip assistance is defined by eight parameters, knee assistance by ten parameters, and ankle assistance by four parameters for a total of 22 parameters. The optimization algorithm can adjust the labeled nodes or state variables (red). The hip stride time is reset at 84% of stride to avoid discontinuities in the desired torque profile during heel strike.

The stride time was calculated as the time between heel strikes averaged over the previous 20 strides. Ankle and knee torque profiles reset at heel strike while the hip torque profile reset at 84% of stride after heel strike. The delayed hip stride timer allowed for torque application during heel strike and prevented discontinuities in the desired torque profile. Timing and duration parameters were defined as a percent of the average stride time.

The exoskeleton accurately applied the desired torque profiles. Torque application was controlled through closed-loop, proportional control with iterative learning [19] and velocity compensation [18]. When no torque was commanded, the device tracked the user’s joint angles with some slack in the system so as not to apply torque. On average, this strategy resulted in 1.6 Nm of root mean square (RMS) error (11% of peak torque). Knee assistance had larger torque tracking error than at the hip or ankle because the state-based periods vary on a step-to-step basis and produce step changes in desired torque. During periods of no torque, the device applied 0.6 Nm of torque on average (Supplementary Section 9).

### 2.3 Optimization Protocol

We first optimized exoskeleton assistance for medium-speed walking, then fast walking and finally slow walking. Participants were most familiar with the medium speed from previous experiments, and we found that transitioning from the fast walking experiment to the slow walking experiment was much easier for the participant than transitioning from slow to fast walking. The medium-speed experiment data for participants 1 and 2 are included in another study [16] which compared single-joint, two-joint and whole-leg assistance strategies.

For each speed, we optimized exoskeleton assistance for at least nine generations, typically over three days. We found this was long enough for the optimization to produce consistent metabolic reductions while maintaining manageable experiment times [16]. On occasion, the optimization took four days because of mid-experiment hardware failures. Participants fasted for two hours before an optimization to minimize the thermal effect of food within a generation and rested at least one day in between experiments. Each generation consisted of 13 conditions. Participants walked in each condition for two minutes, resulting in 26 minutes of walking per generation. With three generations per day, participants walked for at least 78 minutes per optimization experiment. Participants could listen to podcasts while walking with an in-ear, wireless bluetooth headphone.

### 2.4 Optimization Algorithm

We used human-in-the-loop optimization to minimize the metabolic cost of walking. This strategy has been successful to reduce the metabolic cost of walking with exoskeleton assistance [12, 14, 16, 20, 21]. We optimized assistance with the covariance matrix algorithm-evolutionary strategy (CMA-ES), which was successful in previous exoskeleton optimization studies[12, 16, 21]. CMA-ES tests a generation of conditions defined by parameter means and a covariance matrix, ranks the conditions by their performance, then updates the means and covariance matrix of the next generation based on the results of the ranking. We sample each condition for two minutes during which we estimate the metabolic cost through a linear dynamic model [22].

We seeded the optimization with previous, successful profiles. For the first participant, the medium-speed optimization was initialized with the optimized parameters from previous, individual joint optimizations [16]. The optimization for the second participant was seeded based on the optimized parameters from the first participant, and the third participant was initialized with the average of the first two. The fast and slow walking conditions were initialized with the optimized medium-speed parameters for each participant. Torque magnitude parameters were initialized at 75% of the optimized values, and the timing parameters were not changed.

### 2.5 Validation Protocol

We validated the optimized assistance on a separate day after completing the optimization for each speed and before starting the optimization at the next speed. Participants fasted for four hours before validation experiments to eliminate the thermal effect of food over the whole experiment. First, the participant stood quietly for six minutes so we could measure baseline metabolic rate. Then they walked for six minutes in boots, ten minutes in the device without assistance and 20 minutes in the device with optimized assistance. This process was then repeated in reverse to remove ordering effects (ABCDDCBA). Participants were required to rest at least three minutes before walking conditions and at least five minutes before quiet standing so their metabolic rates would return to baseline levels. The conditions were not randomized to minimize the amount of times the participant needed to don or doff the device and to introduce conditions in ascending order of novelty. The increased time for exoskeleton conditions allows the participant’s metabolic rate and kinematic adaptations to stabilize as they adjust to walking in the device.

### 2.6 Measured Outcomes

We measured metabolic rate, muscle activity, ground reaction forces, joint angles, and joint torques during validation experiments. We averaged the outcomes over the last three minutes of each condition to ensure the user’s metabolic rate and gait had reached steady state.

#### 2.6.1 Metabolic Rate

We calculated the metabolic rate through indirect calorimetry using a modified Brockway equation [23] similar to previous studies [12, 16, 21]. Oxygen input and carbon dioxide output were measured with a Cosmed CPET system. Quiet standing measurements were subtracted from the walking condition measurements to calculate the metabolic cost of walking. Measurements were normalized to body mass to allow for comparisons.

Participants wore a cloth or paper mask under the metabolics mask to follow COVID-19 safety protocols for some experiments. Participant 3 wore a cloth or paper mask for all experiments, and participant 2 wore a cloth or paper mask for the slow speed experiments. All other experiments occurred before the pandemic. We found that a cloth or paper mask under the metabolics mask slightly lowers the measured metabolic rate (Supplementary Section 10). This did not affect within-speed comparisons across exoskeleton conditions, nor across-speed comparisons that are normalized within session, for example percent change in metabolic rate.

#### 2.6.2 Muscle Activity

We measured muscle activity with surface electromyography (EMG) (Delsys, Trigno). Gastrocnemius lateralis, vastus lateralis, rectus femoris, semitendinosus and gluteus maximus activity on the participant’s right leg were measured. Measurements were bandpass filtered at 40 and 450 Hz, rectified, then low pass filtered at 10 Hz [24]. We subtracted the minimum signal value and normalized to the peak activity seen in the unassisted condition. Sensor placement was similar to previous biomechanics experiments [25], with some adjustments to avoid interference with the exoskeleton structure and straps. We calculated the RMS of the average stride profiles to evaluate muscle activity changes.

#### 2.6.3 Exoskeleton Torques and Joint Angles

We estimated user kinematics by measuring the exoskeleton joint angles. Magnetic rotary encoders measured the exoskeleton hip, knee and ankle joint angles which provided reasonable approximations of user joint angles. We did not measure kinematics with motion capture because of difficulties with marker occlusion and ghost markers from reflective components on the exoskeleton.

We measured the applied torque at each exoskeleton joint. Strain gauges directly measured applied ankle torque. Load cells measured the applied force at the hips and knees, and we calculated the applied torque for these joints by multiplying the applied force by the exoskeleton lever arm.

#### 2.6.4 Joint Power Calculation

We calculated the exoskeleton joint power by multiplying the joint velocity by the applied torque. We took the average of the positive values to determine the positive power applied. The joint velocity was the time derivative of the joint angles low-pass filtered at 50 Hz. The joint power was summed between the two legs, and the total power was the sum of the joint powers.

#### 2.6.5 Statistical Analysis

We performed two-tailed paired t-tests to determine significance. We compared the assisted conditions to the two control conditions, unassisted and no device, for metabolic cost, cost of transport, stride frequency and RMS muscle activity. We also compared changes from the slow to medium walking speeds, medium to fast walking speeds, and slow to fast walking speeds for positive power changes. For all comparisons, the significance level was *α* = 0.05.

## 3 Results

### 3.1 Metabolic Cost

Exoskeleton assistance reduced the metabolic cost of walking at each speed. Relative to walking with the device turned off (Unasst.), assistance reduced metabolic cost by 26% for slow walking (range 20% to 30%, p = 0.01), 47% for medium-speed walking (range 37% to 53%, p = 0.02), and 50% for fast walking (range 49% to 52%, p = 0.01) (Figure 3 A). This corresponds to a reduction of 0.77 W/kg for slow walking (range 0.60-0.92 W/kg), 1.83 W/kg for medium-speed walking (range 1.54-2.33 W/kg), and 2.65 W/kg for fast walking (range 2.13-3.19 W/kg). Relative to walking with no device (No exo.), assistance reduced metabolic cost by 33% for medium-speed walking (range 21% to 41%, p = 0.04), and 35% for fast walking (range 31% to 39%, p = 0.01). Exoskeleton assistance did not reduce the metabolic cost of slow walking relative to walking with no device.

**Figure 3.**
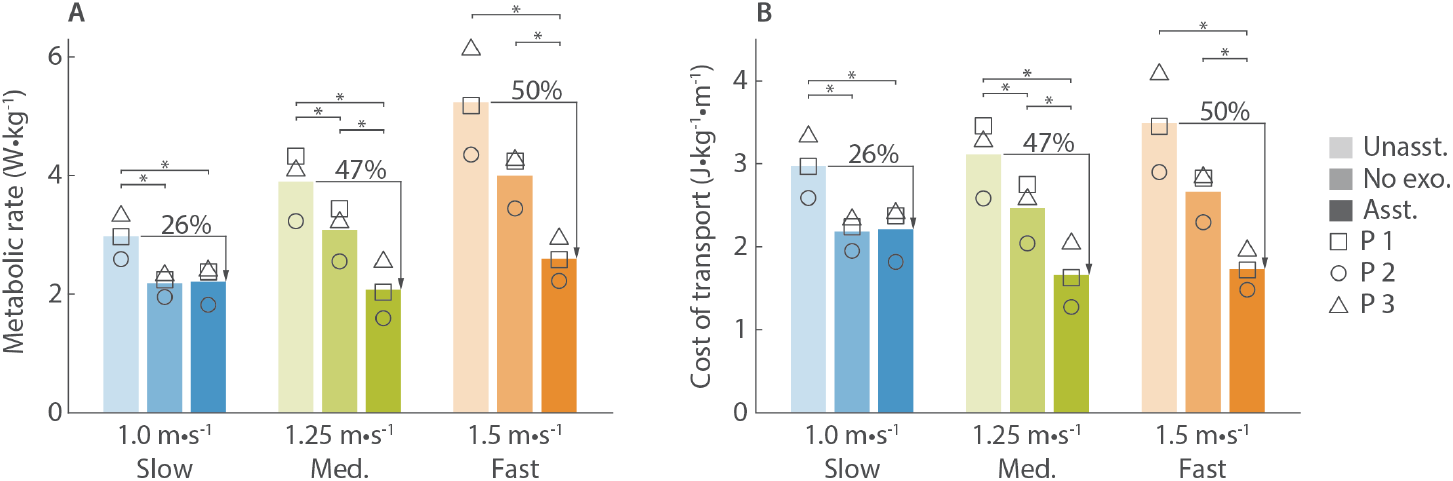
Metabolic cost of walking and cost of transport. (A) The metabolic cost of unassisted walking (Unasst.), walking without the device (No exo.), and assisted walking (Asst.) at slow, medium and fast speeds. Individual metabolic scores are shown with symbols (p1 ☐, p2 O, p3 Δ). Exoskeleton assistance reduced the metabolic cost of walking relative to the unassisted condition at all three speeds and relative to the no device condition for medium and fast walking speeds. Quiet standing has been subtracted from all scores. (B) The cost of transport of unassisted walking, no device walking, and assisted walking at slow, medium and fast speeds. Exoskeleton assistance reduced the cost of transport relative to the unassisted condition at all three speeds. It also reduced the cost of transport relative to the no exoskeleton condition for medium and fast walking speeds.

Exoskeleton assistance decreased the cost of transport relative to walking in the device without assistance. The assisted cost of transport was 2.22 J/kg/m for slow walking (range 1.82-2.44 J/kg/m, p = 0.01), 1.67 J/kg/m for medium-speed walking (range 1.27-2.07 J/kg/m, p = 0.02), and 1.74 J/kg/m for fast walking (range 1.48-1.99 J/kg/m, p = 0.01)(Figure 3 B). The assisted costs of transport for medium and fast walking speeds were similar and smaller than the assisted cost of transport for slow walking.

### 3.2 Joint Power

The total, positive exoskeleton power increased with walking speed (Figure 4 A). The exoskeleton applied 0.73 W/kg for slow walking (range 0.71-0.74 W/kg), 1.08 W/kg for medium-speed walking (range 1.06-1.12 W/kg), and 1.19 W/kg for fast walking (range1.19-1.20 W/kg). Ankle power increased with speed, while there was not a clear trend at the hips or knees (Figure 4 B). Increased metabolic reductions were correlated with increased positive exoskeleton power (Figure 4 C), where a linear fit to the data produced a slope of 3.68 and *R*^2^ = 0.78. The apparent efficiency [26], or the inverse of the slope, was similar to the efficiency of muscles (0.27 and 0.25 respectively).

**Figure 4.**
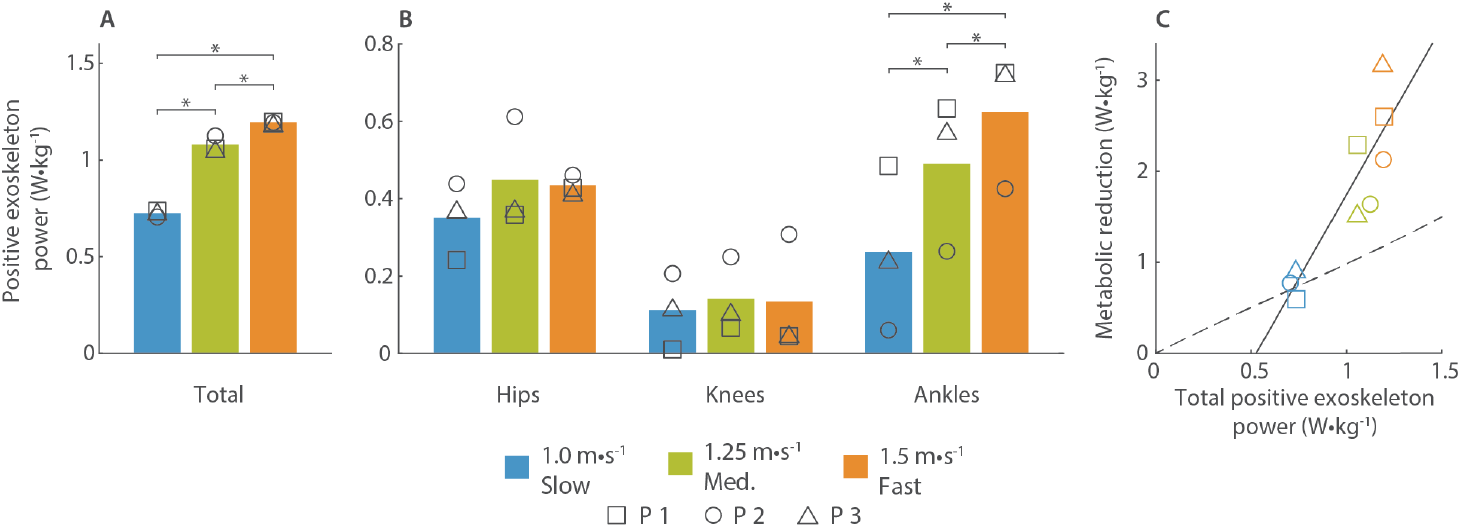
Exoskeleton positive power. (A) total positive exoskeleton power. The positive hip, knee and ankle power was summed across both legs. The positive power is shown for each speed and symbols represent individual participants. (B) positive power at the hips, knees and ankles. The joint power was summed between the left and right legs for each joint. (C) the metabolic rate compared to the positive exoskeleton power. Power applied to each participant is shown with symbols (p1 ☐, p2 O, p3 Δ), and the colors represent walking speeds (blue for slow walking, green for medium-speed walking, and orange for fast walking). The data was fit with a line (solid, 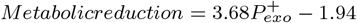, *R*^2^ = 0.78) and unity was shown with a dashed line.

### 3.3 Optimized Assistance

Optimized exoskeleton torque magnitudes varied with speed and participant, but the timing parameters optimized to similar values across speeds and participants (Figure 5). For each participant, ankle plantarflexion torque was the smallest for slow walking, and knee flexion torque was similar for all speeds. Ankle torque for medium and fast walking speeds optimized to the comfort limit (0.80 Nm/kg) in some cases. The torque magnitude changes for hip torque and knee extension torque were participant specific. For example, hip flexion torque increased with speed for participant 1 but was the largest for slow walking for participant 3. Participant 2 saw the largest knee extension torque for fast walking while the other two participants saw similar knee extension torque at all three speeds.

**Figure 5.**
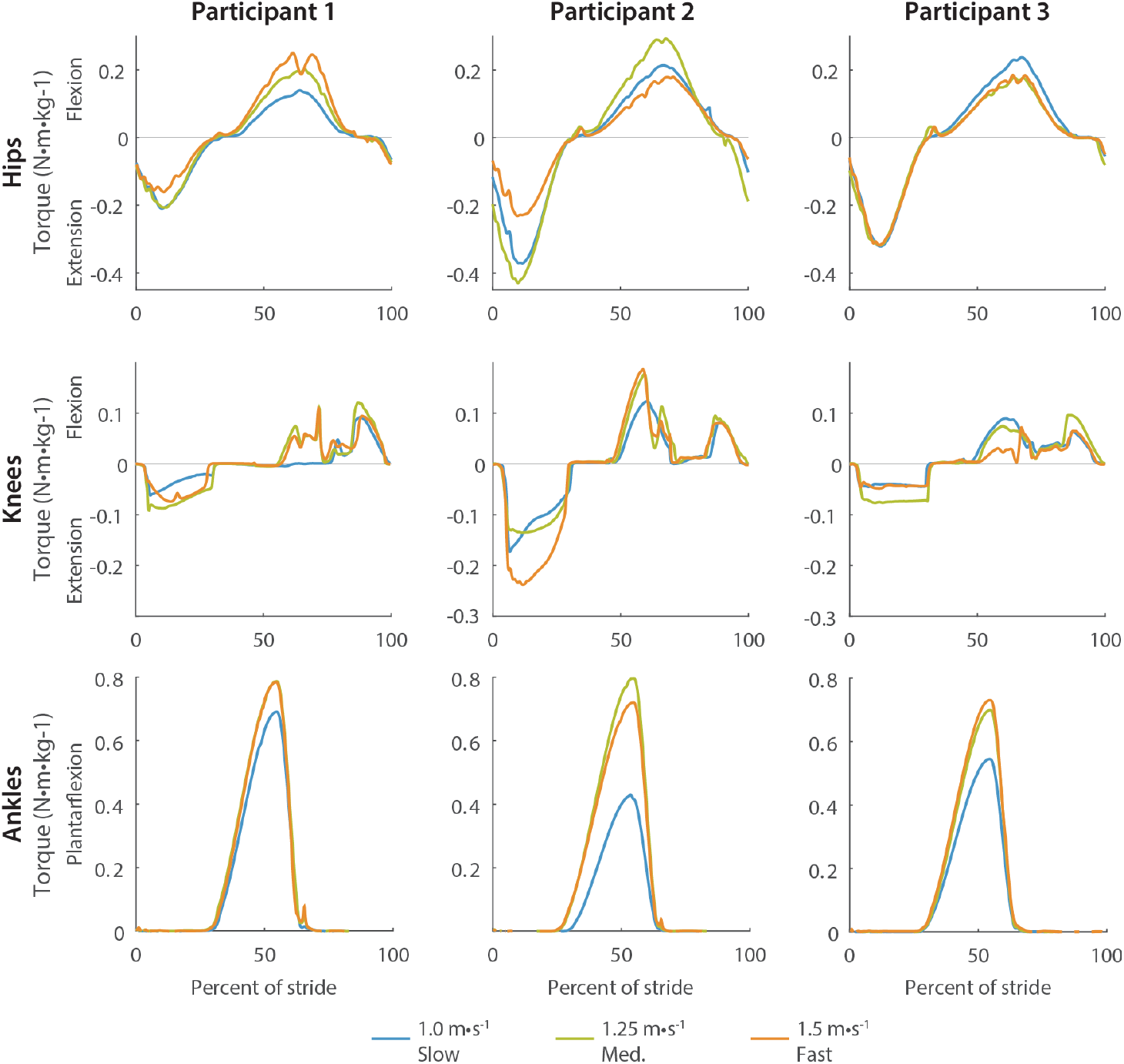
Optimized hip, knee and ankle torque profiles for each participant. (Slow walking (blue) resulted in the smallest ankle torque for all three participants while medium-speed walking (green) and fast walking (orange) resulted in similar ankle torque magnitudes. Timing parameters were similar across speeds and participants.

### 3.4 Kinematics

Exoskeleton assistance resulted in kinematic adaptations in the direction of assistance (Figure 6). For example, at 60% of stride, participants typically plantarflexed their ankles more with assistance than without it and the maximum plantarflexion angle increased with speed. In some instances, assistance exaggerated speed-related joint angle changes. For example, at 50% of stride, the hip extension angle without assistance is typically smaller than medium-speed for slow walking and larger than medium-speed for fast walking. With assistance, the hip extension angle decreased further for slow walking and increased more for fast walking.

**Figure 6.**
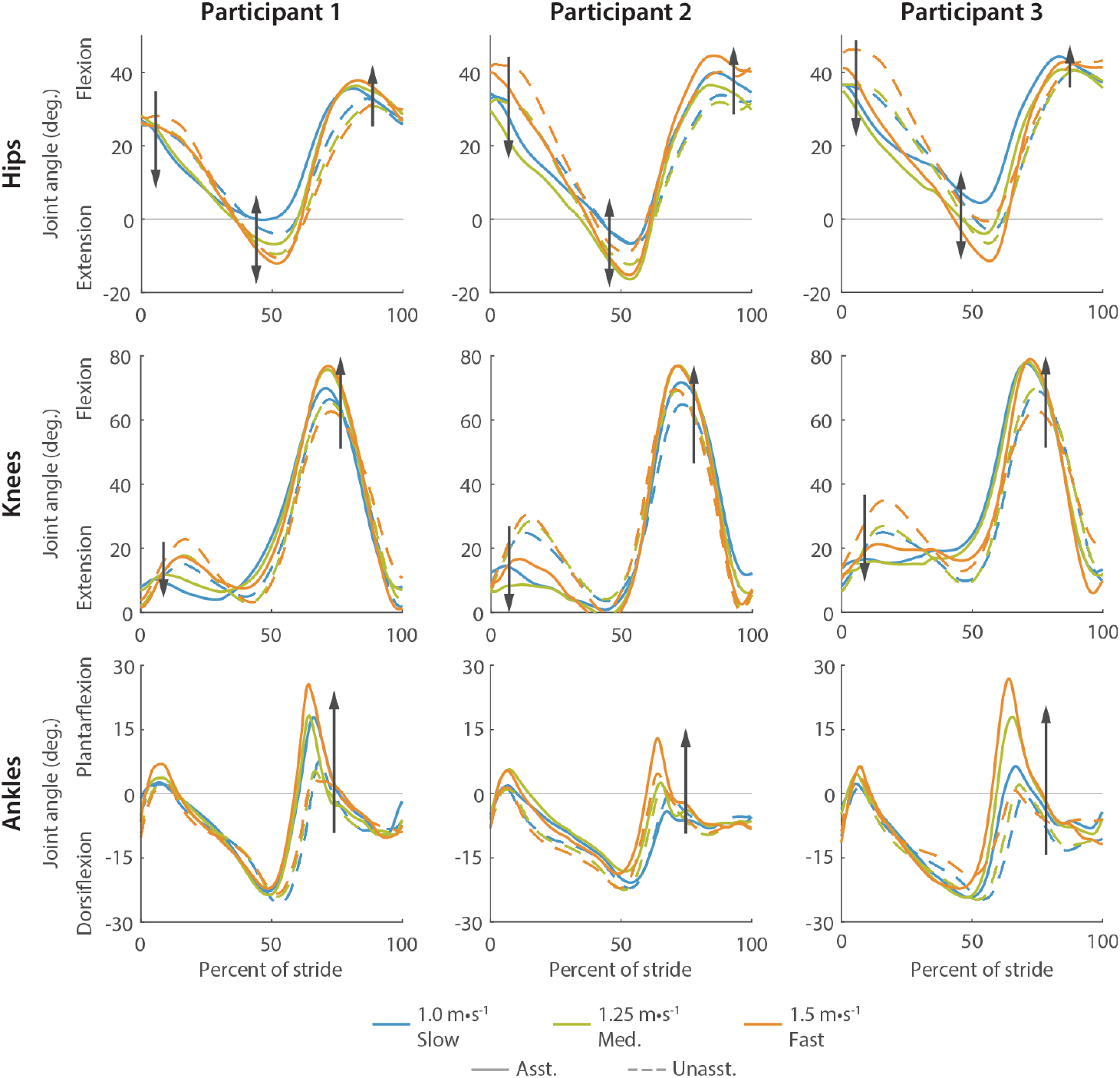
Hip, knee and ankle average joint angle trajectories for each participant. Assisted (solid) and unassisted (dashed) joint angle trajectories are shown for slow (blue), medium-speed (green), and fast (orange) walking. Arrows indicate changes with assistance. For example, the ankle plantarflexion angle increased when assisted at 60% of stride.

The directions of joint angle changes were consistent across participants, but the magnitude of the changes were participant specific. For example, participant 2 plantarflexed their ankles less than participants 1 and 3. Participant 1 had less hip excursion than the other two participants. These participant specific joint angle changes may show different gait strategies, such as a hip strategy or an ankle strategy, or they may be caused by the differences in torque profiles.

Assisted stride frequency was similar at all speeds while the unassisted stride frequency increased with walking speed (Table 1). Participants slightly decreased their stride frequency when walking in the device turned off compared to walking without the device, but this was only statistically significant for medium-speed walking (p = 0.04). Stride frequency increased by 7% (p = 0.005) with assistance relative to unassisted during slow walking and decreased by 4% with assistance relative to unassisted during fast walking (p = 0.06). The stride frequency decrease for fast walking was not statistically significant but was consistent across participants.

**Table 1.**
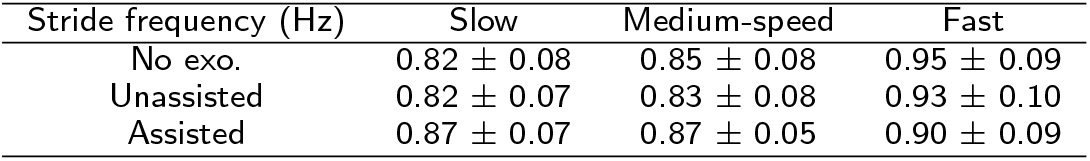
Average stride frequency with no exoskeleton, unassisted and assisted at slow, medium and fast walking speeds.

### 3.5 Muscle Activity

Assistance typically reduced muscle activity relative to the unassisted condition. We normalized the muscle activity to the unassisted condition for each speed, so the maximum unassisted value is always the same. The RMS activity and peak values during assisted walking tended to decrease with speed compared to the unassisted condition (Figure 7), as shown by the gastrocnemius lateralis, rectus femoris and semitendinosus. However, vastus lateralis and gluteus maximus activity did not follow the same trend. Vastus lateralis activity increased around 60% of stride at all three speeds and RMS was always larger than unassisted. This change may indicate a change in coordination strategy that increased vastus energy consumption but reduced energy consumption overall. Gluteus maximus activity did not decrease with assistance for fast walking as it did for medium and slow walking speeds.

**Figure 7.**
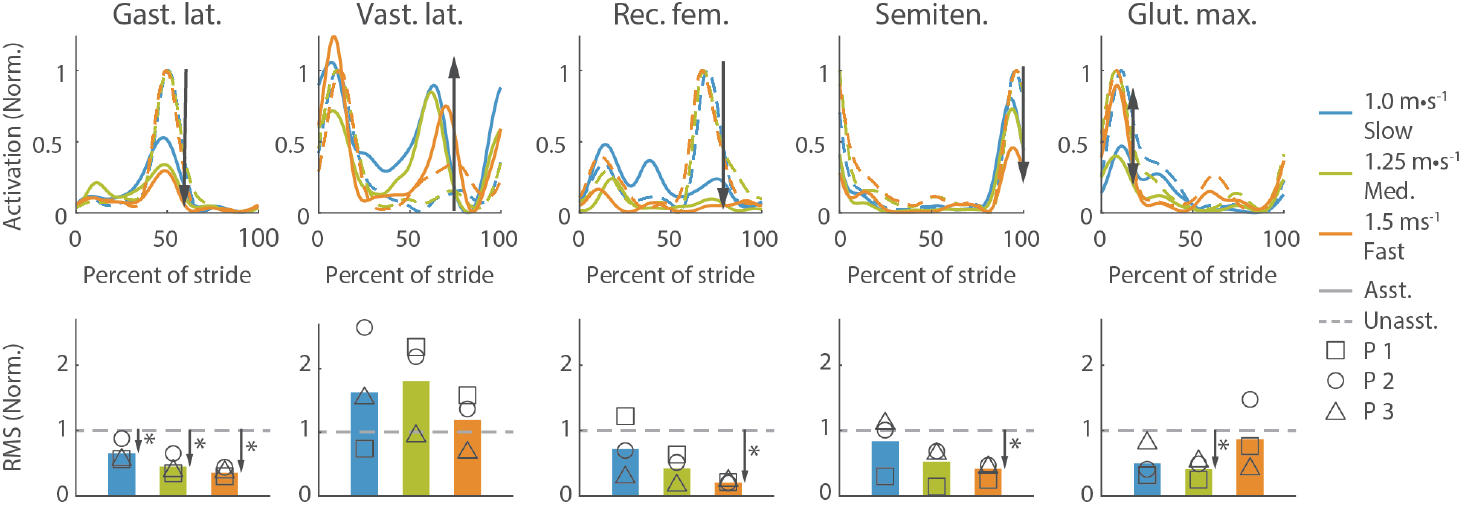
Average muscle activity profile over a stride (top row) and RMS of muscle activity (bottom row). he top row shows the averaged unassisted (dashed) muscle activity profile over a stride and the assisted (solid) for slow (blue), medium-speed (green) and fast (orange) walking. Arrows indicate the muscle activity changes with assistance. The bottom row shows the RMS of the muscle activity with assistance for all three speeds. The RMS of the unassisted muscle activity is shown with the gray line (dashed). Muscle activity was normalized to the unassisted activity resulting in a peak value of 1 for unassisted walking at all speeds.

## 4 Discussion

Exoskeleton assistance can reduce the metabolic cost of walking relative to walking in the device without assistance at slow, medium and fast speeds. Assistance at both medium and fast walking speeds reduced metabolic cost by close to 50%, double the reduction of slow walking, 26%. The absolute reduction in metabolic cost increased with walking speed (0.77 W/kg for slow walking, 1.83 W/kg for medium-speed walking, and 2.65 W/kg for fast walking), similar to our pilot study investigating bilateral ankle assistance at different walking speeds [12]. The percent reductions in metabolic cost for medium and fast walking (47% and 50%) are similar to previous reductions from whole-leg assistance (50%, [16]). These findings demonstrate that exoskeletons can reduce the energy cost of walking at a large range of speeds and are especially effective during fast walking.

Exoskeleton assistance may be more effective at fast speeds than slow speeds because of speed related changes in muscle-tendon mechanics. Elastic energy is stored in tendons while walking, which reduces the needed muscle fiber work and therefore metabolic cost [27, 28]. As walking speed increases, the relative contribution of elastic energy decreases while muscle fiber work increases [29]. Muscle contraction velocity also increases with speed, which increases the amount of metabolic energy required per unit force produced by muscle [30]. Exoskeleton assistance seems to reduce the metabolic cost of walking by replacing muscle fiber work [31]. Since the muscle fibers produce more joint work at fast speeds than slow speeds, the biological work that exoskeleton assistance can replace increases with walking speed. This may be why metabolic reductions increase with speed even without consistent torque trends across participants.

Optimized exoskeleton assistance did not change with speed in the way that biological torque tends to. The optimized torque values were smaller than typical biological torques, ranging from 11% to 55% of biological torque magnitudes. Biological torque typically increases with speed [32] at a larger rate than optimized hip and knee torques. For example, biological knee extension torque increases by 127% when switching from 1.0 to 1.5 m/s [32], while exoskeleton torques increased by only 41% for the participant with the largest change. However, the optimized ankle torque increased by more than biological torque. Biological plantarflexion torque increases by 14% when changing from 1.0 to 1.5 m/s [32], while exoskeleton plantarflexion torque increased for our participants by 36% on average. Some optimized torque values decreased with speed. For example, optimized hip extension torque decreased for fast walking for participants 1 and 2, but biological hip extension torque typically increases with speed. Exoskeleton hip extension torque seems to cause participants to walk with straighter knees [16], so the decreased hip extension torque may have allowed participants to bend their knees more during stance. The optimized torque patterns from this study could inform the design of future exoskeleton products that assist users at a variety of walking speeds.

Participants developed different gait strategies but produced similar trends in total power and metabolic reductions. The optimized torque trends varied on a participant by participant basis, and consequently, the positive power trends at each joint also varied by participant. For example, hip power increased with speed for participant 1, but not for participants 2 and 3. However, the total power was similar for all participants and increased with speed. The standard deviation of the total power was 0.018 W/kg for slow walking (Supplementary Section 5), which is about five times lower than the standard deviation at the individual joints. Trends in metabolic rate were also consistent across participants. The participant specific joint power and torque suggest that participants may have different optimal walking strategies with exoskeleton assistance, and the consistent total power and metabolic reductions suggest that these strategies produce similar outcomes. Exoskeleton assistance may be most effective when customized to the individual, or a wide range of assistance patterns may be similarly effective. To differentiate between these possibilities, future studies could present one participant with optimized assistance patterns from another participant or the average pattern across all participants from this study.

Larger reductions in metabolic rate may be possible with an improved control strategy, frontal plane assistance, or with significantly more exposure to the device. Exoskeleton assistance seems to be unable to reduce the metabolic cost of walking below 2.08 W/kg (Figure 3). The remaining metabolic cost may be necessary for unassisted gait functions like balance or comfort. With changes to exoskeleton hardware and control to better address these aspects of gait, it may be possible to reduce the metabolic cost of walking further. For example, frontal plane assistance incorporating foot placement control might reduce the needed effort for balance. It would have been useful to test improved control strategies that might allow for comfortable application of larger torques. Torque was limited to 0.8 Nm/kg at the ankles for participant comfort, and the optimizer frequently hit this limit, suggesting that higher ankle torque could facilitate greater metabolic cost savings. Finally, wearing the exoskeleton on a daily basis for an extended period of time may lead to greater reductions in metabolic cost as the user may learn to take better advantage of the exoskeleton assistance.

This study could have been improved by testing more participants. Ideally, more participants would complete our protocol, but we were limited due to experiment length (over 40 hours of data collection per participant) and safety concerns with COVID-19. Our tested sample size is large enough to identify statistically significant changes in metabolic rate in part due to the large magnitude of reductions and consistency across the participants [16]. For a desired statistical power of 0.8, a sample size of three participants, and a standard deviation similar to previous exoskeleton optimization studies (7.3% in [12]), the smallest change in metabolic rate we can detect is 24%, which is smaller than the smallest improvement we found here.

The optimized torque profiles identified here may inform the design of future exoskeleton products. Future exoskeletons could target faster walking speeds at which we found exoskeleton assistance to be more effective. The optimized torque patterns we found could define design specifications for exoskeletons that target walking at a variety of speeds. Exoskeleton designers could choose to vary exoskeleton assistance with speed in a similar way to the optimized assistance strategies by increasing ankle plantarflexion torque with speed and applying user-specific changes to hip and knee assistance. Alternatively, there may be a wide variety of effective torque profiles, so exoskeletons may be able to apply the same profiles at all speeds. Future studies could determine the impact of user-specific exoskeleton assistance to further inform exoskeleton assistance changes with speed.

The results of this study could also influence future exoskeleton experiments. Future studies could investigate non-steady state exoskeleton assistance by using the optimized profiles from this study to inform speed-related torque changes. These tests could also compare the efficacy of static assistance to speed-varying and participant-specific assistance. The limited success of assisting slow walking may be a major limitation for exoskeletons intended to assist populations that typically walk more slowly. Optimizing exoskeleton assistance at slower speeds closer to the self-selected speeds of these populations would provide greater insight into this limitation. For such devices, it may be found to be more beneficial to target improvements in population-specific metrics other than metabolic cost, for example increasing self-selected walking speed.

## 5 Conclusions

Exoskeleton assistance can reduce the metabolic cost of walking at a range of speeds, but is most effective for medium and fast walking. Torque changes at the hips and knees varied for each participant, and ankle plantarflexion torque was always smallest for slow walking. While optimized exoskeleton torque varied by participant, the applied power and metabolic reductions were consistent across participants for each speed. This suggests that effective exoskeleton assistance changes with speed may be participant specific, or there may be a wide variety of effective assistance strategies. Exoskeleton products may target faster walking speeds, at which assistance is more effective. The smaller metabolic reductions for slow walking may explain limited success at reducing the metabolic cost of walking for patient populations. It may be more beneficial for exoskeletons to target increasing walking speed for these groups.

## List of Abbreviations

EMG: electromyography
RMS: root mean square

## Ethics approval and consent to participate

All experiments were approved by the Stanford University Institutional Review Board and the US Army Medical Research and Materiel Command (USAMRMC) Office of Research Protections. Participants provided written and informed consent.

## Consent for publication

Participants written and informed consent for publication.

## Availability of data and material

The datasets for this study are available from the corresponding author on reasonable request.

## Author’s contributions

G.B and P.F. designed and constructed the exoskeleton emulator and developed the controllers. G.B., P.F., A.V, S.S. and R.R. conducted the human-subject experiments. G.B., P.F., M.O., K.G. and S.H. designed the experiments. G.B. analyzed the data and drafted and edited the manuscript. P.F. and S.H. also edited the manuscript. S.C. conceived and managed the project and provided design, controls and testing support.

## Acknowledgements

We thank Katie Poggensee for her help with the experimental design, Nick Bianco for his help with controller development, and the Stanford Biomechatronics Laboratory for their feedback and support. This work was supported by the U.S. Army Natick Soldier Research, Development and Engineering Center (Grant number W911QY18C0140).

## References

1. Bornstein, M.H., Bornstein, H.G.: The pace of life. Nature 259(5544), 557–559(1976). doi:10.1038/259557a0. Number: 5544 Publisher: Nature Publishing Group. Accessed 2021-03-24

2. Song, S., Choi, H., Collins, S.H.: Using force data to self-pace an instrumented treadmill and measure self-selected walking speed. Journal of NeuroEngineering and Rehabilitation 17(1), 68 (2020). doi:10.1186/s12984-020-00683-5. Accessed 2021-03-24

3. Nadeau, S., Betschart, M., Bethoux, F.: Gait analysis for poststroke rehabilitation: the relevance of biomechanical analysis and the impact of gait speed. Physical Medicine and Rehabilitation Clinics of North America 24(2), 265–276 (2013). doi:10.1016/j.pmr.2012.11.007

4. Sawicki, G.S., Beck, O.N., Kang, I., Young, A.J.: The exoskeleton expansion: improving walking and running economy. Journal of NeuroEngineering and Rehabilitation 17(1), 25 (2020). doi:10.1186/s12984-020-00663-9. Accessed 2020-07-29

5. Lim, B., Lee, J., Jang, J., Kim, K., Park, Y.J., Seo, K., Shim, Y.: Delayed Output Feedback Control for Gait Assistance With a Robotic Hip Exoskeleton. IEEE Transactions on Robotics 35(4), 1055–1062 (2019). doi:10.1109/TRO.2019.2913318. Conference Name: IEEE Transactions on Robotics

6. Mooney, L.M., Rouse, E.J., Herr, H.M.: Autonomous exoskeleton reduces metabolic cost of human walking during load carriage. Journal of NeuroEngineering and Rehabilitation 11(1), 80 (2014). doi:10.1186/1743-0003-11-80. Accessed 2021-03-24

7. Lee, S., Kim, J., Baker, L., Long, A., Karavas, N., Menard, N., Galiana, I., Walsh, C.J.: Autonomous multi-joint soft exosuit with augmentation-power-based control parameter tuning reduces energy cost of loaded walking. Journal of NeuroEngineering and Rehabilitation 15(1), 66 (2018). doi:10.1186/s12984-018-0410-y. Accessed 2021-03-24

8. Kim, J., Lee, G., Heimgartner, R., Arumukhom Revi, D., Karavas, N., Nathanson, D., Galiana, I., Eckert-Erdheim, A., Murphy, P., Perry, D., Menard, N., Choe, D.K., Malcolm, P., Walsh, C.J.: Reducing the metabolic rate of walking and running with a versatile, portable exosuit. Science 365(6454), 668–672 (2019). doi:10.1126/science.aav7536. Accessed 2021-03-24

9. Malcolm, P., Derave, W., Galle, S., Clercq, D.D.: A Simple Exoskeleton That Assists Plantarflexion Can Reduce the Metabolic Cost of Human Walking. PLOS ONE 8(2), 56137 (2013). doi:10.1371/journal.pone.0056137. Publisher: Public Library of Science. Accessed 2021-03-24

10. Collins, S.H., Wiggin, M.B., Sawicki, G.S.: Reducing the energy cost of human walking using an unpowered exoskeleton. Nature 522(7555), 212–215 (2015). doi:10.1038/nature14288. Number: 7555 Publisher: Nature Publishing Group. Accessed 2021-03-24

11. Seo, K., Lee, J., Lee, Y., Ha, T., Shim, Y.: Fully autonomous hip exoskeleton saves metabolic cost of walking. In: 2016 IEEE International Conference on Robotics and Automation (ICRA), pp. 4628–4635 (2016). doi:10.1109/ICRA.2016.7487663

12. Zhang, J., Fiers, P., Witte, K.A., Jackson, R.W., Poggensee, K.L., Atkeson, C.G., Collins, S.H.: Human-in-the-loop optimization of exoskeleton assistance during walking. Science 356(6344), 1280–1284 (2017). doi:10.1126/science.aal5054. Accessed 2021-03-24

13. Quinlivan, B.T., Lee, S., Malcolm, P., Rossi, D.M., Grimmer, M., Siviy, C., Karavas, N., Wagner, D., Asbeck, A., Galiana, I., Walsh, C.J.: Assistance magnitude versus metabolic cost reductions for a tethered multiarticular soft exosuit. Science Robotics 2(2), 4416 (2017). doi:10.1126/scirobotics.aah4416. Accessed 2021-03-24

14. Ding, Y., Kim, M., Kuindersma, S., Walsh, C.J.: Human-in-the-loop optimization of hip assistance with a soft exosuit during walking. Science Robotics 3(15) (2018). doi:10.1126/scirobotics.aar5438. Publisher: Science Robotics Section: Research Article. Accessed 2021-03-24

15. Cao, W., Chen, C., Hu, H., Fang, K., Wu, X.: Effect of Hip Assistance Modes on Metabolic Cost of Walking With a Soft Exoskeleton. IEEE Transactions on Automation Science and Engineering, 1–11 (2020). doi:10.1109/TASE.2020.3027748. Conference Name: IEEE Transactions on Automation Science and Engineering

16. Franks, P.W., Bryan, G.M., Martin, R.M., Reyes, R., Collins, S.H.: Comparing optimized exoskeleton assistance of the hip, knee, and ankle in single and multi-joint configurations. bioRxiv, 2021–0219431882 (2021). doi:10.1101/2021.02.19.431882. Publisher: Cold Spring Harbor Laboratory Section: New Results. Accessed 2021-03-24

17. Nuckols, R.W., Sawicki, G.S.: Impact of elastic ankle exoskeleton stiffness on neuromechanics and energetics of human walking across multiple speeds. Journal of NeuroEngineering and Rehabilitation 17(1), 75 (2020). doi:10.1186/s12984-020-00703-4. Accessed 2021-03-24

18. Bryan, G.M., Franks, P.W., Klein, S.C., Peuchen, R.J., Collins, S.H.: A hip–knee–ankle exoskeleton emulator for studying gait assistance. The International Journal of Robotics Research, 0278364920961452 (2020). doi:10.1177/0278364920961452. Publisher: SAGE Publications Ltd STM. Accessed 2021-03-24

19. Zhang, C.C. J.and Cheah, Collins, S.H.: Torque control in legged locomotion. In: Sharbafi, S.A. M. (ed.) Bioinspired Legged Locomotion: Models, Concepts, Control and Applications, pp. 347–395. Butterworth-Heinemann, Oxford (2017). Chap. 5. Google-Books-ID: 3gVQCwAAQBAJ

20. Koller, J., Gates, D., Ferris, D., Remy, C.: ‘Body-in-the-Loop’ Optimization of Assistive Robotic Devices: A Validation Study, (2016). doi:10.15607/RSS.2016.XII.007

21. Witte, K.A., Fiers, P., Sheets-Singer, A.L., Collins, S.H.: Improving the energy economy of human running with powered and unpowered ankle exoskeleton assistance. Science Robotics 5(40), 9108 (2020). doi:10.1126/scirobotics.aay9108. Accessed 2021-03-24

22. Selinger, J.C., Donelan, J.M.: Estimating instantaneous energetic cost during non-steady-state gait. Journal of Applied Physiology 117(11), 1406–1415 (2014). doi:10.1152/japplphysiol.00445.2014. Publisher: American Physiological Society. Accessed 2021-03-24

23. Brockway, J.: Derivation of formulae used to calculate energy expenditure in man. Human nutrition. Clinical nutrition 41(6), 463–471 (1987)

24. De Luca, C.J., Donald Gilmore, L., Kuznetsov, M., Roy, S.H.: Filtering the surface EMG signal: Movement artifact and baseline noise contamination. Journal of Biomechanics 43(8), 1573–1579 (2010). doi:10.1016/j.jbiomech.2010.01.027. Accessed 2021-03-24

25. Winter, D.A., Yack, H.J.: EMG profiles during normal human walking: stride-to-stride and inter-subject variability. Electroencephalography and Clinical Neurophysiology 67(5), 402–411 (1987). doi:10.1016/0013-4694(87)90003-4. Accessed 2021-03-24

26. Sawicki, G.S., Ferris, D.P.: Mechanics and energetics of level walking with powered ankle exoskeletons. Journal of Experimental Biology 211(9), 1402–1413 (2008). doi:10.1242/jeb.009241. Publisher: The Company of Biologists Ltd Section: Research Article. Accessed 2021-01-27

27. Cavagna, G.A., Heglund, N.C., Taylor, C.R.: Mechanical work in terrestrial locomotion: two basic mechanisms for minimizing energy expenditure. American Journal of Physiology-Regulatory, Integrative and Comparative Physiology 233(5), 243–261 (1977). doi:10.1152/ajpregu.1977.233.5.R243. Publisher: American Physiological Society. Accessed 2021-03-24

28. Ishikawa, M., Komi, P.V., Grey, M.J., Lepola, V., Bruggemann, G.-P.: Muscle-tendon interaction and elastic energy usage in human walking. Journal of Applied Physiology 99(2), 603–608 (2005). doi:10.1152/japplphysiol.00189.2005. Publisher: American Physiological Society. Accessed 2021-03-24

29. Neptune, R.R., Sasaki, K., Kautz, S.A.: The effect of walking speed on muscle function and mechanical energetics. Gait & Posture 28(1), 135–143 (2008). doi:10.1016/j.gaitpost.2007.11.004. Accessed 2021-03-24

30. Farris, D.J., Sawicki, G.S.: Human medial gastrocnemius force–velocity behavior shifts with locomotion speed and gait. Proceedings of the National Academy of Sciences 109(3), 977–982 (2012). doi:10.1073/pnas.1107972109. Publisher: National Academy of Sciences Section: Biological Sciences. Accessed 2021-03-24

31. Jackson, R.W., Dembia, C.L., Delp, S.L., Collins, S.H.: Muscle–tendon mechanics explain unexpected effects of exoskeleton assistance on metabolic rate during walking. The Journal of Experimental Biology 220(11), 2082–2095 (2017). doi:10.1242/jeb.150011. Accessed 2021-03-24

32. Uchida, T.K., Delp, S.L.: Biomechanics of Movement: The Science of Sports, Robotics, and Rehabilitation. MIT Press, Cambridge, MA (2021). Google-Books-ID: Hu8OEAAAQBAJ

