## Supplementary Material for "Optimized hip-knee-ankle exoskeleton assistance at a range of walking speeds"

**1. Training protocol** 2

**2. Torque parameterization** 3

**3. Metabolic results** 4

**4. Cost of transport** 5

**5. Applied power** 6

**6. Muscle activity profiles** 7

**7. Stride frequency** 8

**8. Parameter ranges and optimized values** 9

Hip parameter ranges and optimized values 9

Knee parameter ranges and optimized values 11

Ankle parameter ranges and optimized values 12

**9. Torque tracking** 13

**10. Impact of cloth mask** 14

### 1. Training protocol

Participants 1 and 2 were considered expert users before beginning the study. They have walked in the device for years developing the control strategy and they completed 12 days (XX hours) of single-joint optimization experiments with this device.

Participant 3 underwent 3 days of training before beginning the optimization experiments. They were gradually exposed to exoskeleton assistance at each joint individually and then all three together. Once all three joints were actuated, we slowly increased the torque magnitudes until they were similar to the optimized results for participants 1 and 2. After 3 days of optimization, we restarted the optimization for participant 3 because we were worried that the optimization algorithm optimized to a local minima. In total, participant 3 completed 5 days of exposure with a variety of assistance strategies.

**Day 1**
On the first training day, the device was fit to the participant. They walked with no torque applied for 15 minutes to become accustomed to walking in the device. They then walked with single joint assistance. They walked for 5 minutes with ankle only torque, 10 minutes with knee only torque, and then 10 minutes with hip only torque. After walking with single joint assistance, we applied hip, knee and ankle torque simultaneously for 10 minutes. We finished the first day of training with 10 minutes of walking with no torque applied. The applied profiles are the average of the first two participant's optimized torque profiles.

**Day 2**
Participant 3 began the second training day with 10 minutes of walking in the device with no torque applied. They then completed three 10 minute periods of simultaneous hip, knee ankle torque. We finished the second day with 10 minutes of walking in the device with no torque applied. The applied torque profiles were the same as the first day of training.

**Day 3**
Participant 3 completed two generations of optimization for the third day of training. We wanted this participant to experience a variety of profiles before beginning the optimization. We also wanted the participant to feel the assistance change every 2 minutes so it would not be jarring during the optimization experiments.

We also spent time making adjustments to the device to address discomfort. This participant felt pressure on their feet, so we changed the insoles to give more space for the participant's foot and alleviate the pressure. We also adjusted padding on the device where it contacts the medial part of the participant's knee.

**Optimization restart**
We restarted the optimization at the medium speed after 9 generations of optimization. We were concerned that this optimization may have been caught in a local minima or that participant 3 had maladapted to the assistance. Exoskeleton assistance visually seemed to alter participant 3's gait more than participant 1 or 2, and the metabolic impact of wearing the device was larger than expected. Participants 1 and 2 saw a 15% increase in metabolic cost when walking in the device without assistance compared to walking without the device. Participant 3 had a 45% increase.

We believe these negative effects were a sign of not enough training, so we restarted the optimization. After restarting the optimization, exoskeleton torques affected participant 3's gait less than before. The metabolic impact of wearing the device also decreased from a 45% increase over walking in boots to a 22% increase suggesting that participant 3 had sufficient training.

#### 2. Torque parameterization

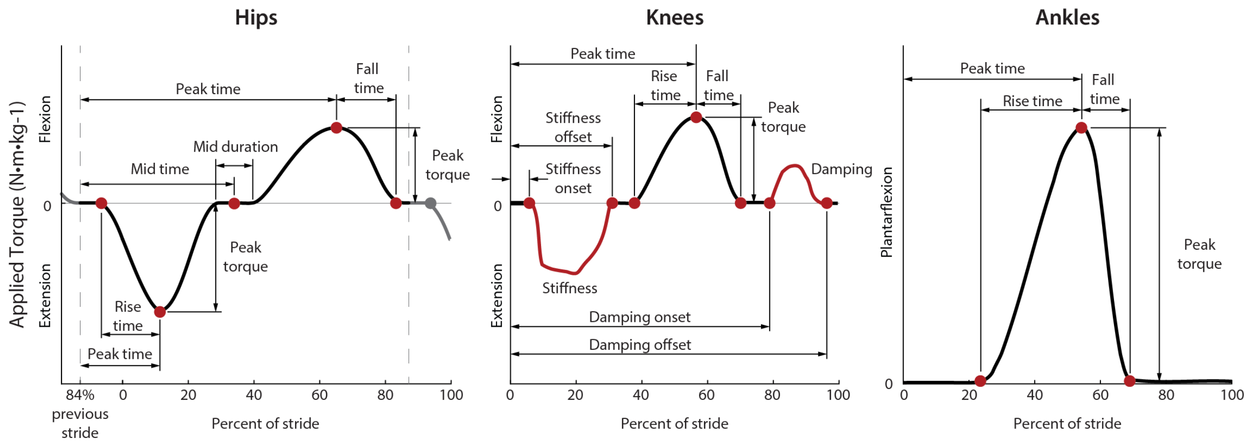


**Figure 1.** Parameterization of the hip, knee and ankle profiles.

The hip profile was defined by 8 parameters (Fig. 2 Hips). These parameters defined the rise time, peak time and peak magnitude of hip extension and the peak time, peak magnitude and fall time of hip flexion. A period of no torque was prescribed between extension and flexion periods and was defined by the mid-point timing and duration.

The knee profile was defined by 10 parameters (Fig. 2 Knees). It consisted of a virtual spring during stance, time based flexion torque near toe off, and a virtual damper during swing. The virtual spring was defined by the stiffness, onset time and offset time. The spring torque was the stiffness multiplied by the knee joint angle, which was set to zero when the knee was straight. Knee flexion torque near toe off was defined by the peak time, peak magnitude, rise time and fall time. The virtual damper during swing was parameterized with a damping coefficient, onset time and offset time, similar to the parameterization of the virtual spring.

The ankle profile was defined by 4 parameters similar to knee flexion near toe-off (Fig. 2 Ankles). These defined the peak time, peak magnitude, rise time, and fall time. Ankle torque was set to zero at 65% of stride at the latest to avoid torque application during swing. This constraint could shorten the fall time, for example a peak time of 55% of stride would result in a maximum fall time of 10% of stride.

### 3. Metabolic results

**Table 1.** Slow walking (1.0 m/s) metabolic cost in W/kg.

| | Participant information | Quiet standing | No exoskeleton | No torque | Optimized torque |
|---|---|---|---|---|---|
| P 1 | 60 kg, 170 cm, F | 1.44 ± 0.14 | 3.72 ± 0.21 | 4.46 ± 0.08 | 3.86 ± 0.04 |
| P 2 | 90 kg, 187 cm, M | 1.50 ± 0.26 | 3.45 ± 0.11 | 4.09 ± 0.02 | 3.32 ± 0.02 |
| P 3 | 80 kg, 182 cm, M | 1.61 ± 0.19 | 3.97 ± 0.43 | 4.97 ± 0.43 | 4.04 ± 0.08 |

**Table 2.** Medium-speed walking (1.25 m/s) metabolic cost in W/kg.

| | Participant information | Quiet standing | No exoskeleton | No torque | Optimized torque |
|---|---|---|---|---|---|
| P 1 | 60 kg, 170 cm, F | 1.77 ± 0.01 | 5.27 ± 0.06 | 6.17 ± 0.12 | 3.84 ± 0.06 |
| P 2 | 90 kg, 187 cm, M | 1.33 ± 0.02 | 3.88 ± 0.09 | 4.56 ± 0.14 | 2.92 ± 0.11 |
| P 3 | 80 kg, 182 cm, M | 1.56 ± 0.02 | 4.82 ± 0.12 | 5.69 ± 0.26 | 4.15 ± 0.07 |

**Table 3.** Fast walking (1.5 m/s) metabolic cost in W/kg.

| | Participant information | Quiet standing | No exoskeleton | No torque | Optimized torque |
|---|---|---|---|---|---|
| P 1 | 60 kg, 170 cm, F | 1.44 ± 0.01 | 5.75 ± 0.29 | 6.71 ± 0.02 | 4.06 ± 0.13 |
| P 2 | 90 kg, 187 cm, M | 1.42 ± 0.09 | 4.86 ± 0.21 | 5.77 ± 0.04 | 3.64 ± 0.02 |
| P 3 | 80 kg, 182 cm, M | 1.73 ± 0.03 | 6.03 ± 0.03 | 7.90 ± 0.01 | 4.71 ± 0.11 |

### 4. Cost of transport

**Table 4.** Slow walking (1.0 m/s) cost of transport in J/kg/m.

| | Participant information | No exoskeleton | No torque | Optimized torque |
|---|---|---|---|---|
| P 1 | 60 kg, 170 cm, F | 2.28 ± 0.06 | 3.02 ± 0.12 | 2.42 ± 0.06 |
| P 2 | 90 kg, 187 cm, M | 1.95 ± 0.09 | 2.59 ± 0.14 | 1.82 ± 0.11 |
| P 3 | 80 kg, 182 cm, M | 2.36 ± 0.12 | 3.36 ± 0.26 | 2.44 ± 0.07 |

**Table 5.** Medium-speed walking (1.25 m/s) cost of transport in J/kg/m.

| | Participant information | No exoskeleton | No torque | Optimized torque |
|---|---|---|---|---|
| P 1 | 60 kg, 170 cm, F | 2.80 ± 0.05 | 3.52 ± 0.10 | 1.66 ± 0.05 |
| P 2 | 90 kg, 187 cm, M | 2.04 ± 0.08 | 2.58 ± 0.11 | 1.27 ± 0.09 |
| P 3 | 80 kg, 182 cm, M | 2.61 ± 0.10 | 3.30 ± 0.20 | 2.07 ± 0.06 |

**Table 6.** Fast walking (1.5 m/s) cost of transport in J/kg/m.

| | Participant information | No exoskeleton | No torque | Optimized torque |
|---|---|---|---|---|
| P 1 | 60 kg, 170 cm, F | 2.88 ± 0.19 | 3.51 ± 0.01 | 1.75 ± 0.09 |
| P 2 | 90 kg, 187 cm, M | 2.30 ± 0.14 | 2.90 ± 0.03 | 1.48 ± 0.02 |
| P 3 | 80 kg, 182 cm, M | 2.87 ± 0.02 | 4.11 ± 0.01 | 1.99 ± 0.07 |

### 5. Applied power

**Table 7.** Positive exoskeleton power in W/kg  at the hips, knees and ankles and the sum of all three.

| | Hips | Knees | Ankles | Total |
|---|---|---|---|---|
| 1.0 m/s | 0.35 ± 0.10 | 0.11 ± 0.10 | 0.26 ± 0.21 | 0.73 ± 0.02 |
| 1.25 m/s | 0.45 ± 0.14 | 0.14 ± 0.10 | 0.49 ± 0.20 | 1.08 ± 0.04 |
| 1.5 m/s | 0.43 ± 0.02 | 0.13 ± 0.15 | 0.62 ± 0.17 | 1.19 ± 0.00 |

**Table 8.** Net exoskeleton power in W/kg at the hips, knees and ankles and the sum of all three.

| | Hips | Knees | Ankles | Total |
|---|---|---|---|---|
| 1.0 m/s | 0.34 ± 0.09 | 0.02 ± 0.11 | 0.18 ± 0.19 | 0.54 ± 0.03 |
| 1.25 m/s | 0.43 ± 0.13 | 0.01 ± 0.11 | 0.37 ± 0.21 | 0.81 ± 0.06 |
| 1.5 m/s | 0.41 ± 0.03 | 0.00 ± 0.16 | 0.53 ± 0.18 | 0.94 ± 0.07 |

**Table 9.** Negative exoskeleton power in W/kg at the hips, knees and ankles and the sum of all three.

| | Hips | Knees | Ankles | Total |
|---|---|---|---|---|
| 1.0 m/s | -0.01 ± 0.01 | -0.09 ± 0.01 | -0.08 ± 0.03 | -0.19 ± 0.03 |
| 1.25 m/s | -0.02 ± 0.01 | -0.13 ± 0.02 | -0.12 ± 0.05 | -0.27 ± 0.05 |
| 1.5 m/s | -0.03 ± 0.01 | -0.13 ± 0.03 | -0.09 ± 0.05 | -0.25 ± 0.08 |

### 6. Muscle activity profiles

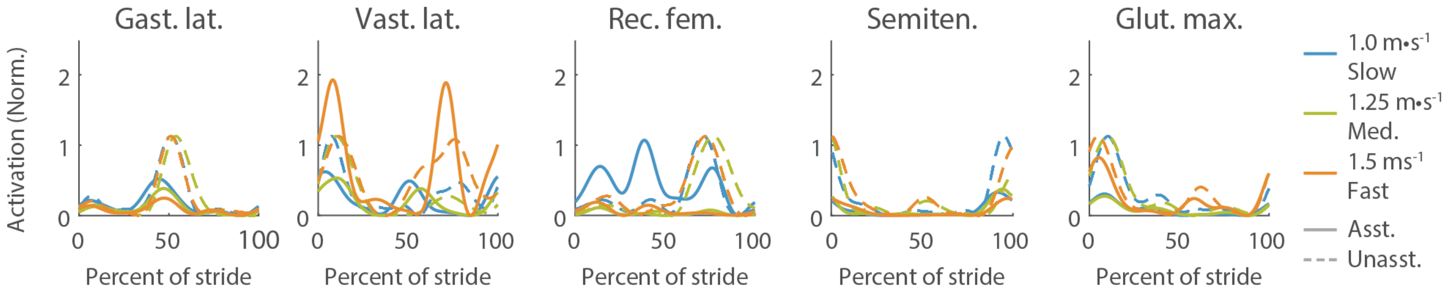


**Figure 2.** Muscle activity averaged over a stride for participant 1.

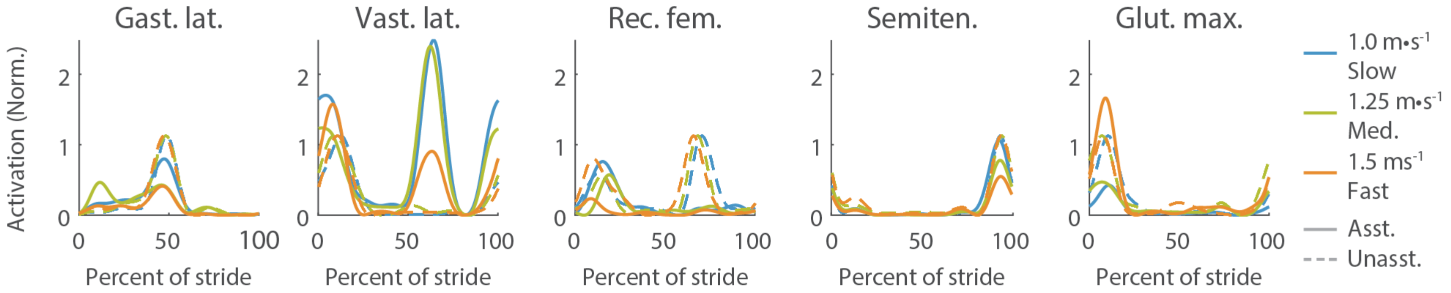


**Figure 3.** Muscle activity averaged over a stride for participant 2.

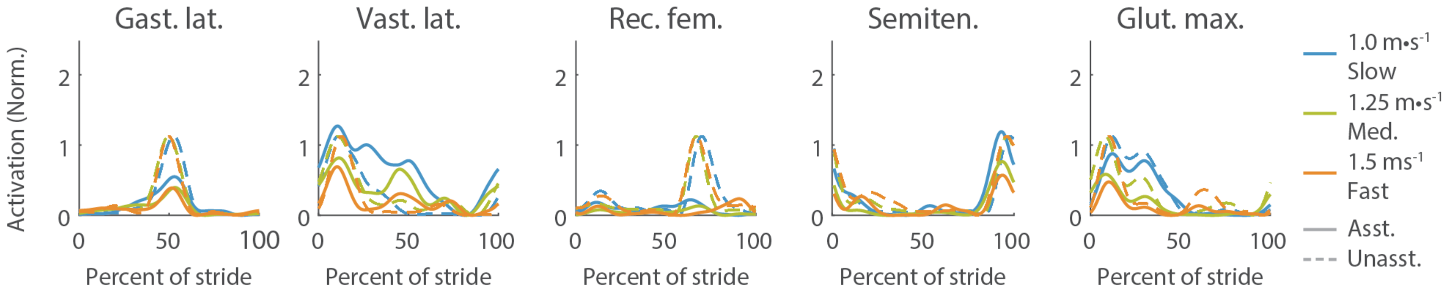


**Figure 4.** Muscle activity averaged over a stride for participant 3.

### 7. Stride frequency

**Table 10.** Stride frequency for participant 1.

| Stride Frequency (Hz) | Slow | Medium-speed | Fast |
|---|---|---|---|
| No exo. | 0.90 ± 0.00 | 0.98 ± 0.00 | 1.05 ± 0.00 |
| Unassisted | 0.90 ± 0.00 | 0.97 ± 0.00 | 1.05 ± 0.00 |
| Assisted | 0.95 ± 0.00 | 0.95 ± 0.00 | 1.00 ± 0.00 |

**Table 11.** Stride frequency for participant 2.

| Stride Frequency (Hz) | Slow | Medium-speed | Fast |
|---|---|---|---|
| No exo. | 0.76 ± 0.00 | 0.83 ± 0.00 | 0.88 ± 0.00 |
| Unassisted | 0.78 ± 0.00 | 0.82 ± 0.00 | 0.87 ± 0.00 |
| Assisted | 0.83 ± 0.00 | 0.85 ± 0.00 | 0.85 ± 0.00 |

**Table 12.** Stride frequency for participant 3.

| Stride Frequency (Hz) | Slow | Medium-speed | Fast |
|---|---|---|---|
| No exo. | 0.78 ± 0.01 | 0.86 ± 0.00 | 0.91 ± 0.00 |
| Unassisted | 0.78 ± 0.00 | 0.85 ± 0.00 | 0.88 ± 0.00 |
| Assisted | 0.84 ± 0.00 | 0.89 ± 0.00 | 0.84 ± 0.00 |

### 8. Parameter ranges and optimized values

The optimization algorithm varied the red nodes and state-based periods. This parameterization was successful in a previous optimization experiment with our device (Franks 2021). We experimentally found the parameter constraints while piloting the assistance. We swept through parameter values to determine the range of comfortable assistance.

The timing parameters are defined as a percent of stride where 1 is 100%. The magnitude parameters are in Nm/kg so we can compare across participants. The medium-speed initial parameters for P2 were based on the optimized assistance at the hip, knee and ankle individually. The initial values for P1 at that speed were based on the optimized values for P2, and the initial values for P3 were the average of the optimized assistance across all participants in Franks 2021.

For slow and fast walking, the initial parameters were based on each participant's optimized parameters from the medium speed. The timing parameters were the same as the medium-speed optimized parameters and the magnitude parameters were set to 75% of the optimized values to allow the assistance and the participant to adapt to the new conditions.

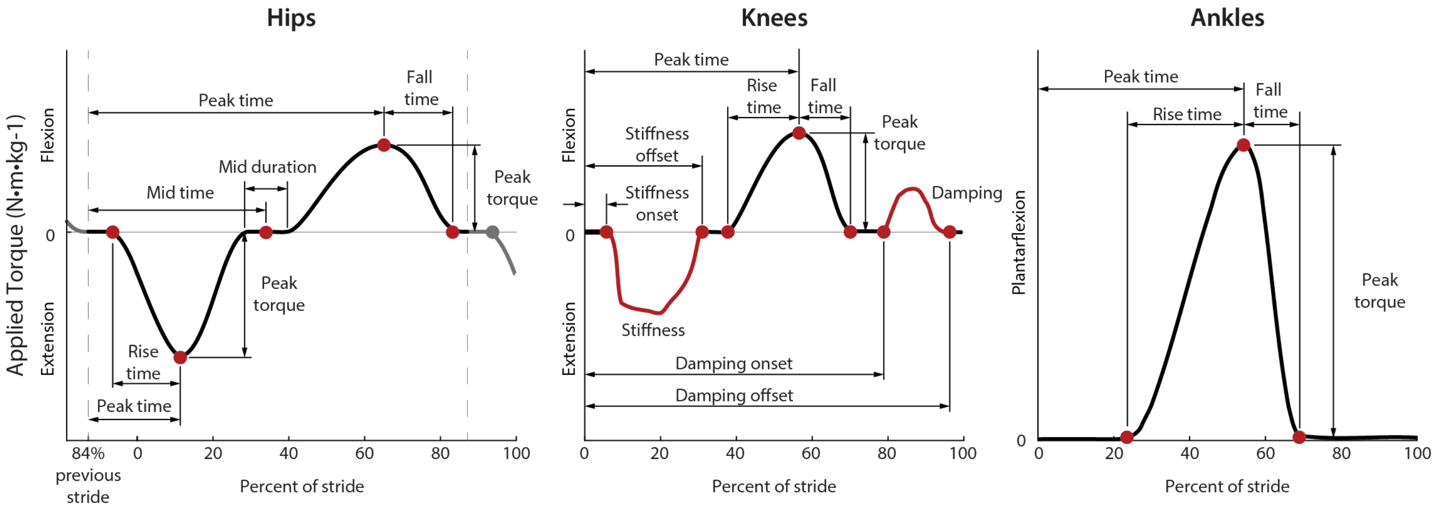


**Figure 5.** Parameterization of the hip, knee and ankle profiles.

#### Hip parameter ranges and optimized values

The hip profile was defined by 8 parameters. It applied hip extension torque through heel strike, so the stride timer began at 84% of stride to avoid discontinuities in the desired profile at heel strike. To convert the hip timing parameters to be based off of heel strike, subtract 16% from the current value.

**Table 10.** Hip parameter ranges for all speeds and initial values for medium-speed walking

| Hips | Hip ext. rise time | Hip ext. peak time | Hip ext. peak torque (Nm/kg) | Mid time | Mid dur. | Hip flex. Peak time | Hip flex. peak torque (Nm/kg) | Hip flex. fall time |
|---|---|---|---|---|---|---|---|---|
| **Min** | 0.125 | 0.225 | 0.000 | 0.450 | 0.000 | 0.750 | 0.000 | 0.125 |
| **Initial P1** | 0.196 | 0.255 | 0.192 | 0.478 | 0.025 | 0.824 | 0.192 | 0.228 |

| | | | | | | | | |
|---|---|---|---|---|---|---|---|---|
| **Initial P2** | 0.176 | 0.258 | 0.420 | 0.472 | 0.020 | 0.814 | 0.271 | 0.244 |
| **Initial P3** | 0.188 | 0.265 | 0.271 | 0.471 | 0.013 | 0.822 | 0.162 | 0.214 |
| **Max** | 0.250 | 0.300 | 0.600 | 0.525 | 0.100 | 0.850 | 0.500 | 0.300 |

**Table 11.** Initial hip parameter values for slow and fast walking

| Hips | Hip ext. rise time | Hip ext. peak time | Hip ext. peak torque (Nm/kg) | Mid time | Mid dur. | Hip flex. Peak time | Hip flex. peak torque (Nm/kg) | Hip flex. fall time |
|---|---|---|---|---|---|---|---|---|
| **Initial P1** | 0.187 | 0.260 | 0.158 | 0.473 | 0.014 | 0.800 | 0.155 | 0.203 |
| **Initial P2** | 0.196 | 0.255 | 0.300 | 0.478 | 0.025 | 0.824 | 0.200 | 0.228 |
| **Initial P3** | 0.178 | 0.266 | 0.245 | 0.468 | 0.008 | 0.808 | 0.165 | 0.220 |

**Table 12.** Optimized hip parameters for slow, medium and fast walking speeds

| 1.0 m/s | HE RT | HE P time | HE torque | Mid time | Mid dur | HF P time | HF torque | HF FT |
|---|---|---|---|---|---|---|---|---|
| **P1** | 0.181 | 0.260 | 0.208 | 0.483 | 0.014 | 0.794 | 0.142 | 0.191 |
| **P2** | 0.211 | 0.268 | 0.373 | 0.484 | 0.029 | 0.831 | 0.209 | 0.259 |
| **P3** | 0.169 | 0.272 | 0.329 | 0.462 | 0.000 | 0.823 | 0.229 | 0.210 |
| **Average** | 0.187 | 0.267 | 0.303 | 0.476 | 0.014 | 0.816 | 0.193 | 0.220 |
| **1.25 m/s** | | | | | | | | |
| **P1** | 0.187 | 0.260 | 0.211 | 0.473 | 0.014 | 0.800 | 0.207 | 0.203 |
| **P2** | 0.196 | 0.255 | 0.425 | 0.478 | 0.025 | 0.824 | 0.283 | 0.228 |
| **P3** | 0.178 | 0.266 | 0.326 | 0.468 | 0.008 | 0.808 | 0.176 | 0.220 |
| **Average** | 0.187 | 0.260 | 0.321 | 0.473 | 0.015 | 0.811 | 0.222 | 0.217 |
| **1.5 m/s** | | | | | | | | |
| **P1** | 0.176 | 0.259 | 0.525 | 0.472 | 0.020 | 0.814 | 0.339 | 0.244 |
| **P2** | 0.196 | 0.250 | 0.160 | 0.474 | 0.006 | 0.813 | 0.272 | 0.188 |
| **P3** | 0.163 | 0.268 | 0.322 | 0.475 | 0.007 | 0.814 | 0.186 | 0.202 |
| **Average** | 0.185 | 0.262 | 0.263 | 0.477 | 0.012 | 0.826 | 0.230 | 0.209 |

#### Knee parameter ranges and optimized values

The knee profile was defined by 10 parameters. The stride time is 0% at heel strike and 100% at the following heel strike of that leg. The knee profile has two state based periods, a virtual spring during stance and a virtual damper during late swing. These two periods were defined by the onset and offset timing of the periods and by the stiffness or damping constant. During the period of virtual spring torque, if the knee joint angle went to 0 before the end of the period, the exoskeleton stopped applying torque for participant comfort.

**Table 13.** Knee parameter ranges for all speeds and initial values for medium-speed walking

| Knees | Stiffness onset | Stiffness *k* | Stiffness offset | Flex. rise time | Flex. peak time | Flex. peak torque | Flex. fall time | Damping onset | Damping coefficient *b* | Damping offset |
|---|---|---|---|---|---|---|---|---|---|---|
| **Min** | 0.001 | 0 | 0.2 | 0.15 | 0.525 | 0 | 0.05 | 0.725 | 0 | 0.9 |
| **Initial P1** | 0.0265 | 0.0073 | 0.2832 | 0.1651 | 0.6105 | 0.1587 | 0.0928 | 0.8074 | 1.1872 | 0.9848 |
| **Initial P2** | 0.025 | 0.008 | 0.271 | 0.205 | 0.588 | 0.247 | 0.094 | 0.812 | 1.763 | 0.968 |
| **Initial P3** | 0.026 | 0.008 | 0.289 | 0.164 | 0.602 | 0.124 | 0.094 | 0.801 | 0.915 | 0.967 |
| **Max** | 0.05 | 0.025 | 0.3 | 0.3 | 0.625 | 0.35 | 0.125 | 0.85 | 2.75 | 0.999 |

**Table 14.** Initial knee parameter values for slow and fast walking

| Knees | Stiffness onset | Stiffness *k* | Stiffness offset | Flex. rise time | Flex. peak time | Flex. peak torque | Flex. fall time | Damping onset | Damping coefficient *b* | Damping offset |
|---|---|---|---|---|---|---|---|---|---|---|
| **Initial P1** | 0.028 | 0.006 | 0.290 | 0.155 | 0.609 | 0.091 | 0.104 | 0.794 | 0.730 | 0.964 |
| **Initial P2** | 0.027 | 0.012 | 0.283 | 0.165 | 0.611 | 0.150 | 0.093 | 0.807 | 1.010 | 0.985 |
| **Initial P3** | 0.024 | 0.004 | 0.300 | 0.170 | 0.600 | 0.064 | 0.099 | 0.782 | 0.980 | 0.975 |

**Table 15.** Optimized knee parameters for slow, medium and fast walking speeds

| 1.0 m/s | KE k on | KE k | KE k Off | KF RT | KF P time | KF T | KF FT | Damp on | Damp *b* | Damp Off |
|---|---|---|---|---|---|---|---|---|---|---|
| **P1** | 0.025 | 0.006 | 0.294 | 0.150 | 0.597 | 0.036 | 0.111 | 0.803 | 0.866 | 0.976 |
| **P2** | 0.025 | 0.012 | 0.295 | 0.150 | 0.603 | 0.135 | 0.086 | 0.816 | 1.359 | 0.956 |
| **P3** | 0.021 | 0.003 | 0.292 | 0.175 | 0.604 | 0.096 | 0.107 | 0.765 | 0.907 | 0.941 |
| **Average** | 0.024 | 0.007 | 0.294 | 0.158 | 0.601 | 0.089 | 0.101 | 0.795 | 1.044 | 0.958 |
| **1.25 m/s** | | | | | | | | | | |
| **P1** | 0.028 | 0.008 | 0.290 | 0.155 | 0.609 | 0.122 | 0.104 | 0.794 | 0.973 | 0.964 |
| **P2** | 0.027 | 0.016 | 0.283 | 0.165 | 0.611 | 0.203 | 0.093 | 0.807 | 1.348 | 0.985 |
| **P3** | 0.024 | 0.005 | 0.300 | 0.170 | 0.600 | 0.085 | 0.099 | 0.782 | 1.307 | 0.975 |
| **Average** | 0.026 | 0.010 | 0.291 | 0.163 | 0.606 | 0.136 | 0.099 | 0.795 | 1.209 | 0.975 |
| **1.5 m/s** | | | | | | | | | | |
| **P1** | 0.022 | 0.004 | 0.272 | 0.150 | 0.607 | 0.114 | 0.102 | 0.805 | 0.636 | 0.966 |
| **P2** | 0.032 | 0.015 | 0.279 | 0.160 | 0.597 | 0.205 | 0.091 | 0.784 | 1.095 | 0.980 |
| **P3** | 0.021 | 0.002 | 0.300 | 0.194 | 0.604 | 0.042 | 0.098 | 0.774 | 0.833 | 0.975 |
| **Average** | 0.025 | 0.007 | 0.283 | 0.168 | 0.603 | 0.120 | 0.097 | 0.788 | 0.854 | 0.974 |

#### Ankle parameter ranges and optimized values

The ankle profile was defined by 4 parameters. The stride time is 0% at heel strike and 100% at the following heel strike of that leg.

**Table 16.** Ankle parameter ranges for all speeds and initial values for medium-speed walking

| Ankles | Peak torque | Peak time | Rise time | Fall time* |
|---|---|---|---|---|
| **Min** | 0.000 | 0.500 | 0.175 | 0.100 |
| **Initial P1** | 0.800 | 0.550 | 0.306 | 0.184 |
| **Initial P2** | 0.800 | 0.550 | 0.400 | 0.200 |
| **Initial P3** | 0.600 | 0.546 | 0.291 | 0.182 |
| **Max** | 0.800 | 0.550 | 0.400 | 0.200 |

*Torque was limited to be applied no later than 65% of stride, so, for example, if peak time was at its latest allowed value (55% of stride), fall time was limited to be 10% of stride.

**Table 17.** Initial ankle parameter values for slow and fast walking

| Ankles | Peak torque | Peak time | Rise time | Fall time* |
|---|---|---|---|---|
| **Initial P1** | 0.600 | 0.550 | 0.282 | 0.173 |
| **Initial P2** | 0.450 | 0.550 | 0.300 | 0.184 |
| **Initial P3** | 0.530 | 0.550 | 0.282 | 0.193 |

**Table 18.** Optimized ankle parameters for slow, medium and fast walking speeds

| 1.0 m/s | Peak torque | Peak time | Rise time | Fall time* |
|---|---|---|---|---|
| **P1** | 0.699 | 0.550 | 0.270 | 0.181 |
| **P2** | 0.428 | 0.542 | 0.261 | 0.200 |
| **P3** | 0.549 | 0.548 | 0.280 | 0.182 |
| **Average** | **0.559** | **0.547** | **0.270** | **0.188** |
| **1.25 m/s** | | | | |
| **P1** | 0.800 | 0.550 | 0.282 | 0.173 |
| **P2** | 0.800 | 0.550 | 0.306 | 0.190 |
| **P3** | 0.707 | 0.550 | 0.282 | 0.193 |
| **Average** | **0.769** | **0.550** | **0.290** | **0.186** |
| **1.5 m/s** | | | | |
| **P1** | 0.800 | 0.548 | 0.273 | 0.172 |
| **P2** | 0.725 | 0.550 | 0.313 | 0.195 |
| **P3** | 0.742 | 0.550 | 0.287 | 0.180 |
| **Average** | **0.756** | **0.549** | **0.291** | **0.182** |

### 9. Torque tracking

**Table 19.** Root mean square torque tracking error for slow, medium, and fast walking speeds.The error is reported in Nm and as percent of the maximum torque.

| 1.0 m/s | Hip | Knee | Ankle |
|---|---|---|---|
| P1 | 0.72 Nm (5.75%) | 1.33 Nm (24.62%) | 1.74 Nm (4.16%) |
| P2 | 0.87 Nm (2.59%) | 3.28 Nm (19.91%) | 0.85 Nm (2.22%) |
| P3 | 0.86 Nm (3.38%) | 1.51 Nm (19.74%) | 1.32 Nm (3.00%) |
| Average | 0.82 Nm (3.90%) | 2.04 Nm (21.42%) | 1.31 Nm (3.12%) |
| 1.25 m/s | | | |
| P1 | 0.79 Nm (6.24%) | 1.88 Nm (24.52%) | 1.61 Nm (3.35%) |
| P2 | 1.13 Nm (2.96%) | 3.48 Nm (19.31%) | 0.90 Nm (1.25%) |
| P3 | 1.17 Nm (4.59%) | 1.95 Nm (22.42%) | 1.72 Nm (3.05%) |
| Average | 1.03 Nm (4.60%) | 2.43 Nm (22.08%) | 1.41 Nm (2.55%) |
| 1.5 m/s | | | |
| P1 | 1.14 Nm (7.01%) | 2.30 Nm (34.50%) | 1.82 Nm (3.80%) |
| P2 | 1.02 Nm (3.72%) | 3.78 Nm (17.51%) | 1.06 Nm (1.62%) |
| P3 | 1.02 Nm (3.99%) | 1.59 Nm (25.49%) | 1.46 Nm (2.43%) |
| Average | 1.06 Nm (4.91%) | 2.56 Nm (25.83%) | 1.45 Nm (2.62%) |

### 10. Impact of cloth mask

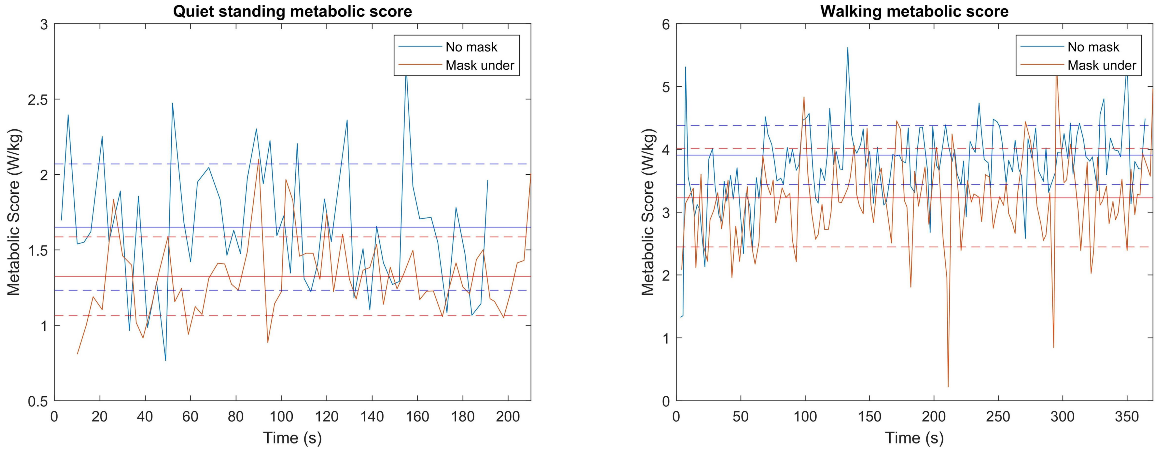


**Figure 6.** Metabolic impact of the cloth mask under the metabolic mask. Participant 1 measured their metabolic cost of quiet standing and walking on the treadmill for 6 minutes each. All measurements were collected in the same session. They measured their metabolic cost with no cloth mask (blue) and with a cloth mask under the metabolics mask (red). The average of the last 3 minutes of walking is shown with the solid line, and the dashed lines show the range for one standard deviation. The participant was the only person in the lab space, so they did not put others at risk by not wearing a mask. The cloth mask lowered the metabolic cost by 0.33 W/kg for the standing condition and by 0.68 W/kg for the walking condition.
